# Systems-wide analysis of the GATC-binding nucleoid-associated protein Gbn and its impact on *Streptomyces* development

**DOI:** 10.1101/2021.02.06.430045

**Authors:** Chao Du, Joost Willemse, Amanda M. Erkelens, Victor J. Carrion, Remus T. Dame, Gilles P. van Wezel

## Abstract

Bacterial chromosome structure is organized by a diverse group of proteins collectively referred to as nucleoid-associated proteins (NAPs). Many NAPs have been well studied in Streptomyces, including Lsr2, HupA, HupS, and sIHF. Here, we show that SCO1839 represents a novel family of Actinobacteria NAPs and recognizes a consensus sequence consisting of GATC followed by (A/T)T. The protein was designated Gbn for <u>G</u>ATC-<u>b</u>inding <u>N</u>AP. Deletion of *gbn* led to alterations in development and antibiotic production in *Streptomyces coelicolor*. Chromatin immunoprecipitation sequencing (ChIP-Seq) detected more than 2800 binding regions, encompassing some 3600 GATCWT motifs, which comprise 55% of all such motifs in the *S. coelicolor* genome. DNA binding of Gbn in vitro increased DNA stiffness but not compaction, suggesting a role in regulation rather than chromosome organization. Transcriptomics analysis showed that Gbn binding generally leads to reduced gene expression. The DNA binding profiles were nearly identical between vegetative and aerial growth. Exceptions are SCO1311 and SCOt32, for a tRNA editing enzyme and a tRNA that recognises the rare leucine codon CUA, respectively, which nearly exclusively bound during vegetative growth. Taken together, our data show that Gbn is a highly pleiotropic NAP that impacts growth and development in streptomycetes.

**IMPORTANCE:** A large part of the chemical space of bioactive natural products is derived from Actinobacteria. Many of the biosynthetic gene clusters for these compounds are cryptic, in others words, they are expressed in nature but not in the laboratory. Understanding the global regulatory networks that control gene expression is key to the development of approaches to activate this biosynthetic potential. Chromosome structure has a major impact on the control of gene expression. In bacteria, the organization of chromosome structure is mediated by a diverse group of proteins referred to collectively as nucleoid-associated proteins (NAPs), which play an important role in the control of gene expression, nucleoid structure and DNA repair. We here present the discovery of a novel and extremely pleiotropic NAP, which we refer to as Gbn. Gbn is a sporulation-specific protein that occurs only in the Actinobacteria and binds to GATC sequences, with a subtle but broad effect on global gene expression. The discovery of Gbn is a new step towards better understanding of how gene expression and chromosome structure is governed in antibiotic-producing streptomycetes.

## INTRODUCTION

Streptomycetes are filamentous soil bacteria with a complex life cycle, which are well known for their ability to produce various kinds of antibiotics and other valuable natural products. Thus, they are a major source of clinical drugs (1–3). The life cycle of Streptomyces starts with the germination of a spore that grows out to form vegetative hyphae. Exponential growth is achieved via tip extension and branching, eventually resulting in a dense mycelial network (1, 4). When the environmental situation requires sporulation, for example due to nutrient starvation, streptomycetes start their reproductive growth phase by developing aerial hyphae, which eventually differentiate into chains of unigenomic spores (5, 6). The production of antibiotics temporally correlates with the onset of development (7, 8). The complexity of the underlying regulatory networks is underlined by the fact that the

*Streptomyces coelicolor* genome encodes some 900 regulatory proteins, of which only a minute fraction has been functionally characterized (9). Many of these affect the control of development and antibiotic production, such as the bld and whi genes that are responsible for the control of aerial hyphae formation and sporulation, respectively, and global regulatory genes such as adpA, afsR, dasR and atrA that pleiotropically control antibiotic production (10).

The control of chromosome structure is an important factor in the control of gene expression. In bacteria, the organization of chromosome structure is mediated by a diverse group of proteins referred to collectively as nucleoid-associated proteins (NAPs) (11–13). These are generally small DNA binding proteins involved in processes such as controlling gene expression, nucleoid structure, or DNA repair. Well-known NAPs in Streptomyces include Lsr2, HupA, HupS, sIHF, and IHF. Lsr2 binds non-specifically to AT-rich sequences and can globally repress gene expression (14). HupA and HupS are homologs of HU (for histone-like protein from strain U93) proteins, which are differentially regulated depending on the developmental growth phase (15). sIHF is one of the basic architectural elements conserved in many actinobacteria and is able to influence the regulation of secondary metabolism and cell development (16). IHF binds a well conserved nucleotide sequence, while HU binds to random DNA sequences (17), yet with a preference for bent, distorted or flexible DNA (18). A proteomic survey of *Streptomyces coelicolor* identified 24 proteins with NAP-like properties, namely the known Lsr2, HupA, HupS and sIHF and 20 yet unidentified proteins (19). Although the functions of many NAPs are still not clear, BldC for example has a major impact on the transcriptome (20, 21).

We previously showed via pull-down assays that the candidate NAP SCO1839 binds to the promoter region of the cell division regulatory gene ssgR (22). SsgR is the transcriptional activator of the cell division activator gene ssgA (23). SsgA and its paralogue SsgB are both required for sporulation (24–26), and together coordinate the onset of sporulation-specific cell division in Streptomyces, whereby SsgB directly recruits the cell division scaffold protein FtsZ to the future sites of septation (27).

In this study, we show that SCO1839 represents a novel family of small DNA binding proteins, which plays a role in the regulation of Streptomyces development and antibiotic production. The protein is specific to the Actinobacteria, with an HTH DNA binding motif containing three helices, and plays a role in the control of morphogenesis. Chromatin immunoprecipitation coupled with massive parallel DNA sequencing (ChIP-Seq) revealed that the protein binds to over 2800 genomic regions with one or more binding sites, recognising a specific DNA binding motif centred around the consensus sequence GATC. Thus, we designated the protein Gbn for <u>G</u>ATC-<u>b</u>inding <u>N</u>AP. Transcriptomics data showed that genes bound by Gbn on the promoter regions tends to express more in *gbn* mutant, suggesting a suppressive effect of Gbn.

## MATERIALS AND METHODS

### Reagents

All restriction enzymes are ordered from NEB (Massachusetts, U.S.), including BamHI-HF (Cat. R3136), XbaI (R0145), EcoRI-HF (R3101), HindIII-HF (R3104), BbsI (R0539), NcoI-HF (R3193), SnaBI (R0130), StuI (R0187), SacI (R0156), NdeI (R0111). Phusion polymerase (M0532) and T4 DNA ligase (M0202) were also obtained from NEB. Plasmid mini-prep kit (Cat. 740727.250) was from BIOKE (Leiden, The Netherlands). DNA purification was achieved using DNA Clean & Concentrator kit (Cat. D4029) from Zymo Research (California, U.S.).

### Biological Resources

#### Strains and growth conditions

All strains used in this study are listed in Table S3. Escherichia coli strain JM109 was used for routine cloning, E. coli ET12567 (28) for preparing non-methylated DNA, ET12567 containing driver plasmid pUZ8002 (29) was used in conjugation experiments for introducing DNA to Streptomyces. E. coli strains were grown in Luria broth at 37°C supplemented with the appropriate antibiotics (ampicillin, apramycin, kanamycin and/or chloramphenicol at 100, 50, 25 and 25 μg·mL^-1^ respectively) depending on the vector used. *S. coelicolor* A3(2) M145 was the parent strain for all mutants. Streptomyces strains were grown on soya flour medium (SFM) for conjugation, SFM agar medium or MM agar medium supplemented with 0.5% mannitol for phenotype characterization, R5 agar plates for protoplast regeneration, and MM agar medium supplemented with 0.5% mannitol covered with cellophane for ChIP-Seq and transcriptomics culture growth. Solid cultures were grown in a 30°C incubator unless described specifically. For liquid cultures, approximately 10^6^ spores were inoculated in 100 mL Erlenmeyer flask equipped with steel spring containing 15 mL TSBS (tryptone soya broth sucrose) medium (30). The flasks were incubated at 30°C with constant shaking at 180 rpm. Antibiotics used for screening Streptomyces transformants were apramycin and thiostrepton (20 and 10 μg·mL^-1^, respectively).

#### Constructs and cloning

Primers used for PCR and short double strand DNA fragment are listed in Table S4. PCR was preformed using Phusion DNA polymerase using standard protocol as described previously (31). All plasmids and constructs described in this study are summarized in Table S5. The constructs generated in this study were verified by sanger sequencing performed in BaseClear (Leiden, The Netherlands).

The *gbn* knock-out strategy was based on the unstable multi-copy vector pWHM3 as described previously (32). Briefly, up- and down-stream region of *gbn* were amplified from genome and cloned into pWHM3. Between these two regions, an apramycin resistance cassette from pGWS728 (33) was inserted as selection marker. The resulting vector pGWS1255 is a knock-out construct that can replace nucleotide positions the +1 to +207 of *gbn* with apramycin resistance cassette, where +1 refers to the translation start site. The apramycin resistance cassette was subsequently removed using Cre expressing construct pUWL-Cre (34, 35) yielding the clean knock-out strain GAD003. For complementation of the *gbn* deletion mutant, the nucleotide positions −565 to +228 relative *gbn* translation start site, containing entire coding region of *gbn* (with stop codon) and its promoting region was cloned into low copy number plasmid pHJL401, yielding complementation construct pGWS1260. This construct was then transformed to GAD003, resulting in strain GAD014.

To perform ChIP-Seq experiment, the 3×FLAG was fused to the end of original copy of *gbn* on genome using codon optimised CRISPR-Cas9 system (36). The spacer sequence located at the end of gbn and was inserted into the pCRISPomyces-2 plasmid as described by (36). Template for homology-directed repair (HDR) which ensures the insertion of 3×FLAG sequence was cloned into pCRISPomyces-2 followed by spacer insertion (Figure S3), yielding *gbn*-3×FLAG knock-in construct pGWS1298. Mutagenesis was done according to (36). A successful 3×FLAG tag knock-in strain was identified by PCR and sequencing, designated GAD043.

For over-expression of *gbn*, the ermE promoter was cloned to replace part of the original promoter region of *gbn* using the same CRISPR-Cas9 system. A spacer sequence were designed at the promoter region of *gbn* and this was inserted into pCRISPomyces-2 (36). HDR template was designed to remove −157 to +4 region of *gbn* and replace this region with PermE sequence yielding pGWS1295. PermE was digested from pHM10a (37). Following the same procedure as above, strain GAD039 was obtained which expresses *gbn* from the ermE promoter.

To produce His_6_-Gbn for electrophoretic mobility shift assay (EMSA) experiments, *gbn* was cloned into protein expression construct pET28a and transformed into E. coli strain BL21 CodonPlus (DE3)-RIPL (Invitrogen, Massachusetts, U.S.). To generate methylated and non-methylated DNA for EMSA, part of the promoter region of *gbn* and a random region that is not bound by Gbn were cloned into pUC19 (38). This yielded vector pGWS1300 that contains a Gbn binding domain and pGWS1451 contains a stretch of DNA that is not bound by Gbn.

### Database mining and clustering

The protein sequence of Gbn (SCO1839) from *S. coelicolor* was used as query in HMMER web server (39) to obtain all Gbn-like proteins from the database, resulting in 727 hits. Sequences with an E value < 0.01 (684 sequences) were selected to generate a Hidden Markov model (HMM) profile using HMMER suit v3.1b2 (40). This profile was used to search against a custom database containing 146,856 genomes with all available bacteria genomes (access date Feb. 9, 2019). Hits with E-value ≤ 5.5 × 10^-9^ (2,317 sequences) were aligned to the generated HMM profile using the hmmalign tool from HMMER suit. Using the alignment, a network was built calculating the pairwise distance between all the detected Gbn proteins and the threshold for clustering was settled at 0.8. Network visualizations were constructed using Cytoscape v3.7.1 (41).

### Time-lapse confluent plate morphology monitoring

Approximately 10^7^ spores were plated on MM agar supplemented with mannitol. The plates were then placed upside down in Perfection V370 scanner (Epson, Nagano, Japan) located inside 30°C incubator. A scanning picture was taken every hour, and images were processed using custom python script to get the brightness value of the plate. Specifically, the pictures were first converted to grey scale. 70% the diameter of the plate from the centre was selected as the region of interest (ROI). The average grey value of all the pixels within ROI was used as the brightness of the mycelium lawn. The measured values from one plate were then normalized to range 0 to 1.

### Scanning electron microscopy (SEM)

Mycelia were grown on MM agar supplemented with mannitol and grown for 5 days. Sample preparation and imaging was done as described before (42, 43), using JSM-7600F scanning electron microscope (JEOL, Tokyo, Japan). For each strain, 5 images with 7,500 × magnification were taken in randomly selected spore-rich areas. The length and width of spores in each picture were measured using ImageJ version 1.52p strictly according to a randomized file list, to minimize selection bias. Only spores which are approximately parallel to the focal plane were measured.

### DNA-protein cross-linking and chromatin immunoprecipitation

10^8^ spores of strain GAD043 were plated on MM agar medium specified above. After 25 h or 48 h growth, cellophane disks were soaked up-side-down in PBS solution containing 1% formaldehyde for 20 min allowing DNA-protein crosslink. Ten plates were collected for 25 h samples; four plates were collected for 48 h samples. Then the disks were moved to PBS solution containing 0.5 M glycine for 5 min to stop crosslinking reaction. The mycelium was then collected, washed in PBS and resuspended in 0.5 mL lysis buffer (10 mM Tris-HCl pH 8.0, 50 mM NaCl, 15 mg·mL^-1^ lysozyme, 1× protease inhibitor (Roche, Bavaria, Germany) and incubated at 37°C for 20 min. After incubation, 0.5 mL IP buffer (100 mM Tris-HCl pH 8.0, 250 mM NaCl, 0.8% v/v Triton-X-100) was added to the sample, and chromosomal DNA sheared to 100 to 500 bp using Bioruptor Plus water bath sonication system (Diagenode, Liège, Belgium). After centrifugation to remove cell debris, lysates were incubated with 40 μL Anti-FLAG M2 affinity gel (cat A2220, Sigma-Aldrich, St. Louis, U.S.) suspension according to the manufacturer’s instructions and incubated at 4°C overnight. After centrifugation and washing, the pellets and 50 μL of untreated total extracts (controls) were incubated in 100 μL IP elution buffer (50 mM Tris-HCl pH7.5, 10 mM EDTA, 1% m/v SDS) at 65°C overnight to reverse cross-links. Beads were then removed by centrifugation before DNA extraction with phenol-chloroform. The DNA sample was then extracted with chloroform and the water layer was further purified using DNA Clean & Concentrator kit (Cat. D4029, Zymo Research, California, U.S.). The samples were then sent for next generation sequencing using the BGI-Seq platform (BGI, Hong Kong, China).

### ChIP-Seq data analysis

Clean reads received from the sequencing contractor were aligned to the *S. coelicolor* M145 genome with RefSeq accession number NC_003888.3 using bowtie2 v2.34 (44). Resulted SAM files are sorted using SAMtools v1.9 (45) producing BAM files. MACS2 v2.1.2 (46) was then used for binding peak prediction and peak modelling by comparing the chromatin immunoprecipitated DNA sample with the corresponding whole genome sample. The models for both samples are shown in Figure S2. The enrichment data used in Figure 3 was calculated for each nucleotide using MACS2 ‘bdgcmp’ command with ‘-m FE’ switch. The peak summit positions including sub peak positions of each predicted binding region were then extracted, and the ± 150 bp region of each summit was extracted from the genome sequence using a python script dependent on the Biopython module v1.70 (47). Extracted sequences were subjected to MEME-ChIP v5.02 (48), which is suitable for large sequence sets, for binding motif prediction.

**Figure 3.**
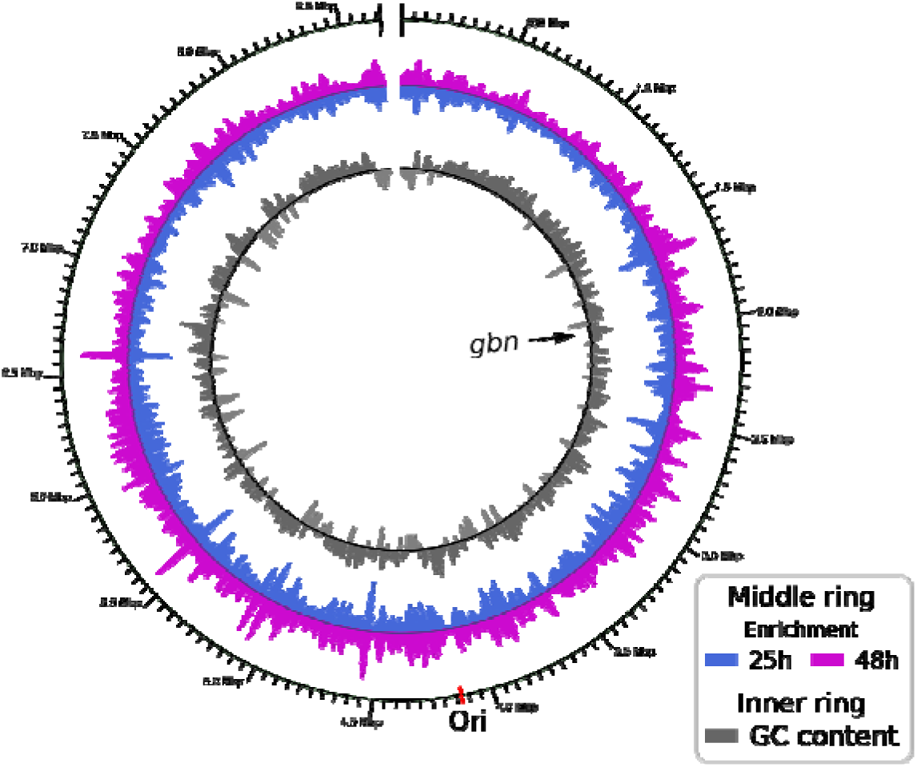
Genome-wide distribution of Gbn protein binding sites along the *S. coelicolor* genome. The outer ring shows the genome location; the middle ring shows the local average fold enrichment from ChIP-Seq analysis, 48 h sample oriented outwards, 25 h sample oriented inwards; the inner ring shows the local average G+C content. The G+C percentage above median is plotted outwards, below median inwards. Bin size for local averaging was 20,000 bp. *gbn* location have been indicated by black arrow.

The enrichment data of two samples was averaged separately in a moving bin of 20,000 bp and plotted in opposite directions as the middle ring of the circular genome diagram. The G+C content was calculated using the same moving bin and centred at the middle of maximum and minimum value before plotting as the inner ring on the plot (Figure 3). For determining the overlap of low G+C content regions and high enrichment regions, the genome was divided into 1,000 bp long sections, the G+C content and average enrichment levels were calculated. The sections with G+C content below the first quartile was considered low in G+C, and those with average enrichment level above the third quartile were considered high in enrichment. To find genes possibly regulated by Gbn, the locations of promoter regions (−350 to +50) of all genes were extracted from the genome containing annotations and checked for overlap with ± 150 bp location of the summit of Gbn binding peaks. This was done using a python script dependent on the module Biopython and pybedtools v0.8 (49) and external BEDTools v2.27 (50).

### Transcriptomics and data analysis

Spores (10^8^ cfu) of *S. coelicolor* M145 and *gbn* knockout strain GAD003 were plated on MM agar plates overlayed with cellophane disks. After 24 h (vegetative growth) or 45 h (aerial growth) of growth, mycelia were scraped from the cellophane disks, snap frozen in liquid N_2_ and disrupted using TissueLyser (Qiagen, Venlo, The Netherlands) for 3 times 30 s at 30 Hz. Total RNA extracted using the Kirby method described (51). The RNA-seq libraries preparation and sequencing were outsourced to Novogene (Novogene Europe, Cambridge, UK). Ribosomal RNA was removed from the samples using NEBNext® Ultra™ Directional RNA Library Prep Kit (NEB, Massachusetts, U.S.). Sequencing libraries were generated using NEBNext® Ultra™ RNA Library Prep Kit for Illumina® (NEB, Massachusetts, U.S.) and sequencing was carried on an Illumina® NovaSeq™ 6000 platform. Raw data was cleaned using fastp v0.12.2 (52), then mapped to the *S. coelicolor* M145 genome (GenBank accession AL645882.2) using bowtie2 v2.4.4 (53). Read counts for each gene were generated using featureCounts v2.0.1 (54). Transcripts per million (TPM) values were generated using custom python script, differently expressed genes and log_2_ fold change (LFC) was generated using DESeq2 v1.32.0 (55) with the data shrinkage function “apeglm” (56).

### Electrophoretic mobility shift assay (EMSA)

Gbn-His_6_ was expressed and purified as described previously (57). Purified protein was dialyzed over-night at 4°C against EMSA buffer (10 mM Tris-HCl, pH 7.9, 0.1 mM EDTA, 50 mM KCl). 50 bp double strand DNA was generated by gradual cooling of reverse complemented single strand oligonucleotides in T4 DNA ligase buffer (NEB, Massachusetts, U.S.) from 95°C to 12°C in 45 min. pGWS1300 was extracted from DAM methylation effective E. coli strain JM109 and DAM deficient *E. coli* strain ET12567, while pGWS1451 for negative control was extracted from strain ET12567 only. The target fragments were then digested and blunted using DNA polymerase I Klenow fragment (NEB, Massachusetts, U.S.). The *in-vitro* DNA-protein interaction studies were done in EMSA buffer in a total reaction volume of 10 μL, the reactions were incubated at 30°C for 15 min. The whole reaction volume was then loaded on 5% polyacrylamide gels and separated by electrophoresis. The gel was briefly stained with ethidium bromide and imaged in a Gel Doc imaging system (BioRad, California, U.S.).

### Tethered particle motion

Tethered particle motion experiments were carried out as described previously (58) with minor modifications. The experimental buffer used was 10 mM Tris-HCl pH 8.0, 0.1 mM EDTA, 50 mM KCl, 0.5% acetylated BSA. Data was collected for each protein concentration at least in duplicate. An anisotropic ratio cut-off of 1.3 and a standard deviation cut-off of 8% were used to select single-tethered beads. The region of interest (−609 to +33 bp relative to the *gbn* translation start site) was amplified from the genome and then inserted into pUC19 using Gibson assembly yielding plasmid pGWS1462. DNA was then amplified as a 685 bp fragment from this construct using a forward primer labelled with biotin (Biotin-CTGGCTGAAACGGAATAGGT) and a reverse primer labelled with digoxygenin (Digoxygenin-AGCTCAGCGAGAACCGG).

## RESULTS AND DISCUSSION

### SCO1839 is a small NAP specific to Actinobacteria

We previously identified SCO1839 as a DNA binding protein that binds to the promoter region of the sporulation regulatory gene ssgR (22). SsgR activates transcription of ssgA, which encodes a pleiotropic developmental regulator and activator of sporulation-specific cell division in streptomycetes. SCO1839 is a small protein of 73 amino acids (7.6 kDa) with a predicted isoelectric point (pI) of 10.53, indicative of an alkalic protein. A Pfam sequence search (59) did not yield any significant matches to known protein families, suggesting that SCO1839 is the first member of a novel protein family. It was suggested that SCO1839 may be a nucleoid-associated protein (NAP) (19).

To obtain more insights into the distribution and phylogeny of SCO1839, a conserved Hidden Markov Model (HMM) domain was constructed using all SCO1839-like proteins from Streptomyces species. Consequently, an HMM search against all available bacteria full genomes in the database was performed. No hits were found outside the order Actinomycetales, strongly suggesting that SCO1839 is an Actinobacteria-specific protein (Figure 1A and B). Eight main clusters of similar groups of homologs were found. The largest cluster mainly consists of SCO1839 orthologs from Streptomyces, Amycolatopsis, Pseudonocardia, Frankia, and Actinomadura. Other major clusters include a cluster of Nocardia and Rhodococcus, Micromonospora and Salinispora, Geodermatophilus and Blastococcus. Orthologs from Rhodococcus form two separate clusters. Interestingly, 24 Actinomadura species and 38 Streptomyces species have two paralogues of SCO1839, which divide into two additional clusters. For most genera, more than 90% of the species encode at least one copy of SCO1839-like proteins. The genus Rhodococcus forms an exception, as only 28.7% (92 out of 321) of the sequenced genomes of this genus contain a SCO1839-family protein (Figure 1A), and these proteins divided into three distinct clusters. This could be related to the fact that it is the only genus among all listed genera that does not have a true mycelial life style (1). The low conservation in the non-sporulating *Rhodococcus* suggests that SCO1839 may primarily be sporulation specific.

**Figure 1.**
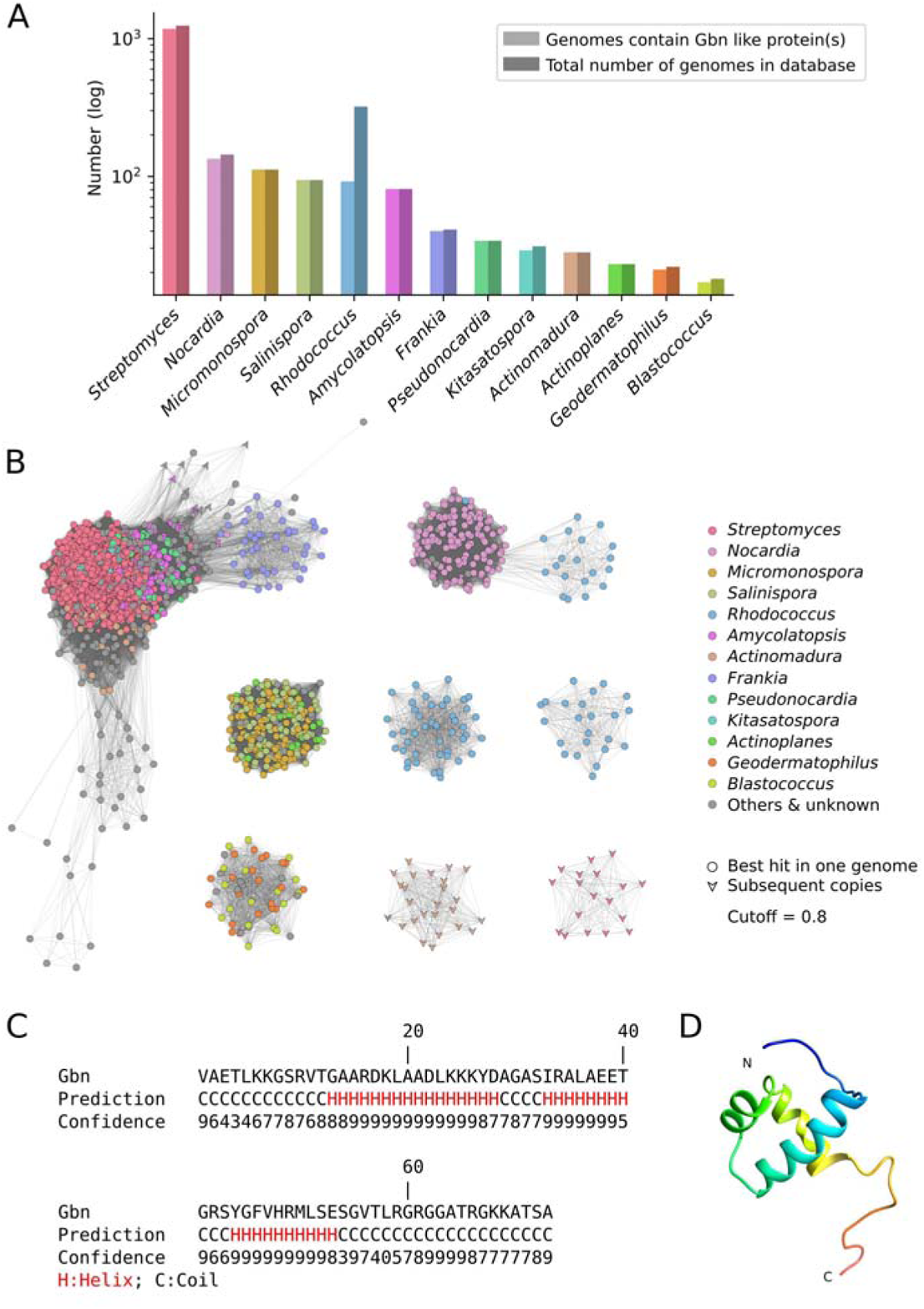
Distribution, phylogenetic network and structural analysis of Gbn-family proteins. **(A)** Bar plot representing the distribution of Gbn-like proteins in all genomes used in this study. Light colours indicate the number of sequenced genomes containing Gbn-like proteins in specific genera, dark colours represent the total number of sequenced genomes in each genus. **(B)** Sequence similarity network built with a threshold of 0.8 using all Gbn-like proteins detected in this study. Nodes represent Gbn-like proteins. Edges connecting the nodes represent phylogenetic distances. Colours indicate the taxonomic affiliation of each orthologue. Shapes denote whether Gbn is the primary copy in one genome. **(C)** Predicted secondary structure of Gbn, obtained from the I-TASSER output. **(D)** *In silico* structural model of the Gbn protein generated with I-TASSER.

*In silico* structural modelling of SCO1839 was performed using the I-TASSER server (60), revealing a putative single DNA binding helix-turn-helix (HTH) motif, in the form of a tri-helical structure (Figure 1C and 1D). No homology was seen to any other known transcriptional regulator family from bacteria. Very short aa stretches flank the residues belonging to the HTH motif. Nine other protein structures found in PDB (Protein data bank, rcsb.org, 61) share structural analogy to SCO1839, and have similar HTH motifs. These are proteins found in a wide range of organisms and with different functions, including a DNA helicase from the archaeon *Pyrococcus furiosus* (PDB structure ID: 2ZJ8), *E. coli* (2VA8, 2P6R) and from humans (5AGA), a ribosomal protein from yeast (5MRC), a human cell division cycle protein (2DIN), a yeast terminator binding protein (5EYB), a regulator from Staphylococcus aureus (2R0Q), and a tRNA synthetase from the archaeon *Archaeoglobus fulgidus* (2ZTG). However, none of these proteins were as small as SCO1839 and neither contained only one DNA binding motif. To the best of our knowledge, no other bacterial protein with similar structure has been reported before. Therefore, we propose that the SCO1839-like proteins form a new family of bacterial DNA binding proteins. SCO1839 is further nominated Gbn, for <u>G</u>ATC-<u>b</u>inding <u>N</u>AP (see below).

### Gbn binds to thousands of DNA binding sites with GATC as core motif

To obtain insights into the genome-wide DNA binding capacity of Gbn, ChIP-Seq analysis was performed on samples harvested after 25 h (vegetative growth) and 48 h (sporulation). Following this approach all binding sites of Gbn on the *S. coelicolor* chromosome can potentially be identified. For this purpose, the original copy of *gbn* on the genome was fused with a sequence encoding a triple FLAG tag at its 5’-terminus using CRISPR-Cas9 (see Materials and Methods section). The strain had a phenotype that was highly similar to that of the parent (Figure S1). The ChIP-Seq data showed a wide distribution of Gbn binding events (Figure 3). Half of the binding sites with a strong enrichment (50% of the sites at 25 and 38 h) colocalized with low G+C content regions. In total, 2825 and 2919 binding regions were identified using MACS2 software (46) for samples obtained from mycelia in the vegetative and sporulation stage, respectively. Interestingly, there was a near complete overlap (> 90%, 2402) between the Gbn binding events found in the two samples (Pearson correlation coefficient 0.945, Figure 3). This not only shows that the binding specificity of Gbn is largely growth phase-independent, but also that the experiments were highly reproducible. The result also indicates that while the expression of *gbn* is higher during sporulation, the specificity of the protein for its binding sites does not change, as shown by the highly similar binding profiles from the ChIP-seq experiments.

To obtain a consensus binding site for Gbn, the binding regions from ChIP-Seq results were extracted and modelled using MACS2 and MEME-ChIP (46, 48). In this way, the sequences GATCAT and GATCTT were identified as binding sites, and thus GATCWT represents the Gbn binding site (Figure 4A). The most conserved binding core was GATC, which is a palindrome known as recognition site for DNA methylation (62). The predicted motifs also showed G/C preference on the flanking region separated by gaps of two base pairs (Figure 4A). It is important to note that virtually all binding regions that were identified as significant (> 99.8%) contained a GATC motif, and most (88.2% for 25 h, 84.0% for 48 h) contained the consensus sequence GATCWT. The *S. coelicolor* genome contains in total 6,501 GATCWT sequences, 54.5% and 55.5% were present in the predicted binding regions in the 25 h and 48 h samples, respectively. Note that many binding sites have more than one copy of the motif. Closer inspection of the raw data of the ChIP-seq data revealed that in fact all GATCWT motifs on the *S. coelicolor* genome are covered by an increased sequencing coverage, but not all of them can be reliably detected by MACS2 without increasing the false positives to an unacceptable level. This was not seen for GATC sequences not followed by AT or TT. We believe that Gbn can bind most GATCWT sequences on the *S. coelicolor* chromosome. However, not all GATCWT sequences may be accessible to Gbn either directly because they are occupied by other proteins or indirectly because of chromosome organization of the region. Further investigation of this phenomenon is needed.

**Figure 4.**
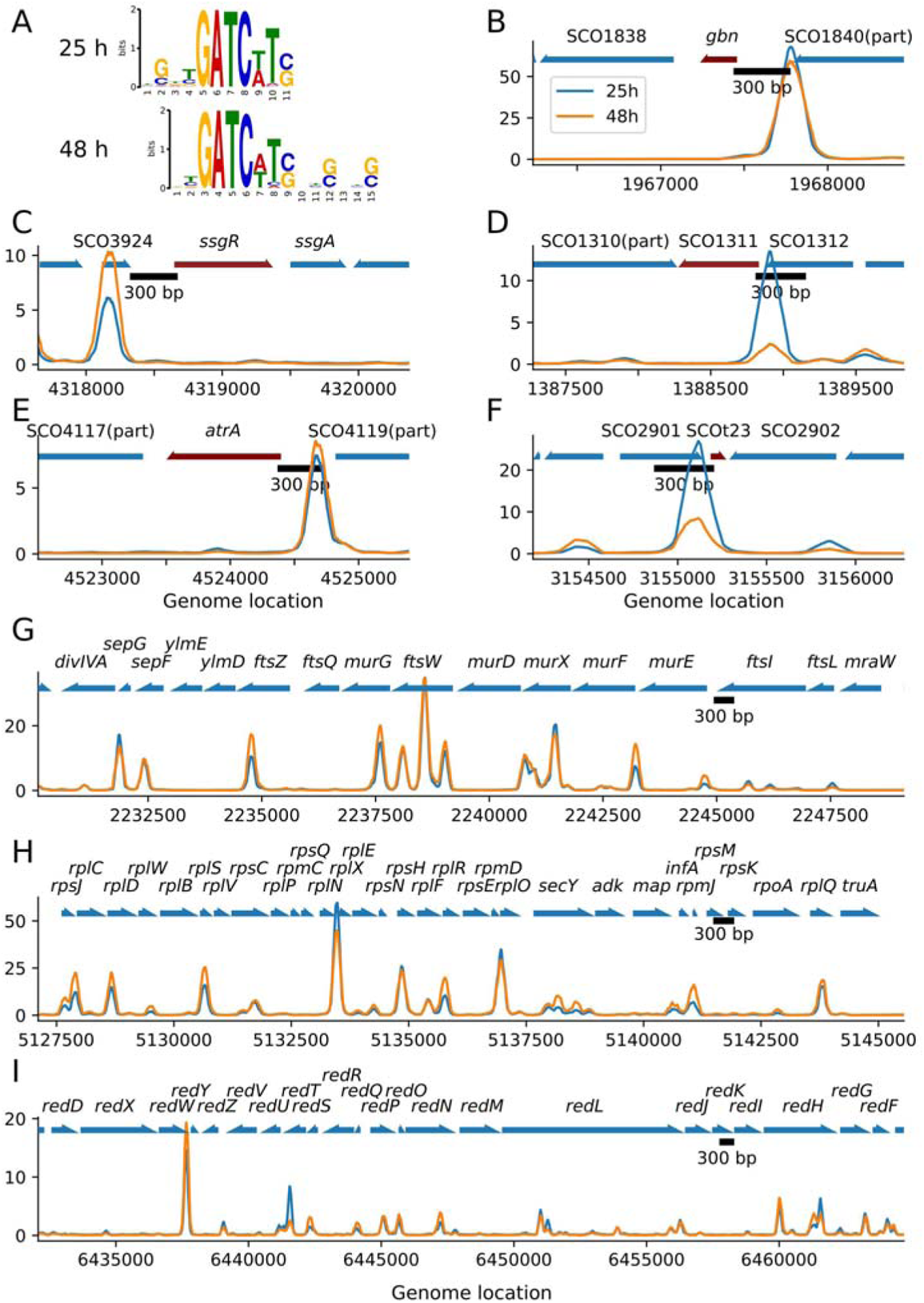
Gbn protein binding sites analysis. **(A)** Gbn DNA binding motifs predicted by combining the programs MACS2 and MEME-ChIP. (B-I) Enrichment level (Y axis) of Gbn in target region as measured by ChIP-Seq experiment, corresponding samples of blue and orange lines are indicated by the legend in B. B-F, targeted genes with flanking regions (1000 bp) are included. Red arrow indicating target gene, other genes are coloured blue. G-I, *dcw* gene cluster, ribosomal protein gene cluster, and *red* gene cluster, respectively. Note that the y-axes of the plots are in automatic scale.

The affinity of Gbn for its DNA binding motif was further examined in vitro using electrophoretic mobility shift assays (EMSA). Result show that Gbn could bind to the GATC motif in vitro. The additional nucleotide (AT)T on one side increased the affinity of Gbn, with the half binding concentration reduced by more than a third (Figure 5B). Furthermore, when more GATC motifs were present in the DNA fragments, simultaneous binding of more Gbn proteins was observed (Figure 5C and D). For the short 50 bp DNA fragment with four target DNA motifs, only three binding events can be observed from EMSA experiment, possibly only one of the two GATC sequences in close proximity could be bound by one Gbn protein as a consequence of steric hindrance. Taken together, Gbn showed good binding to a motif centred around GATC in experiments *in vitro*.

**Figure 5.**
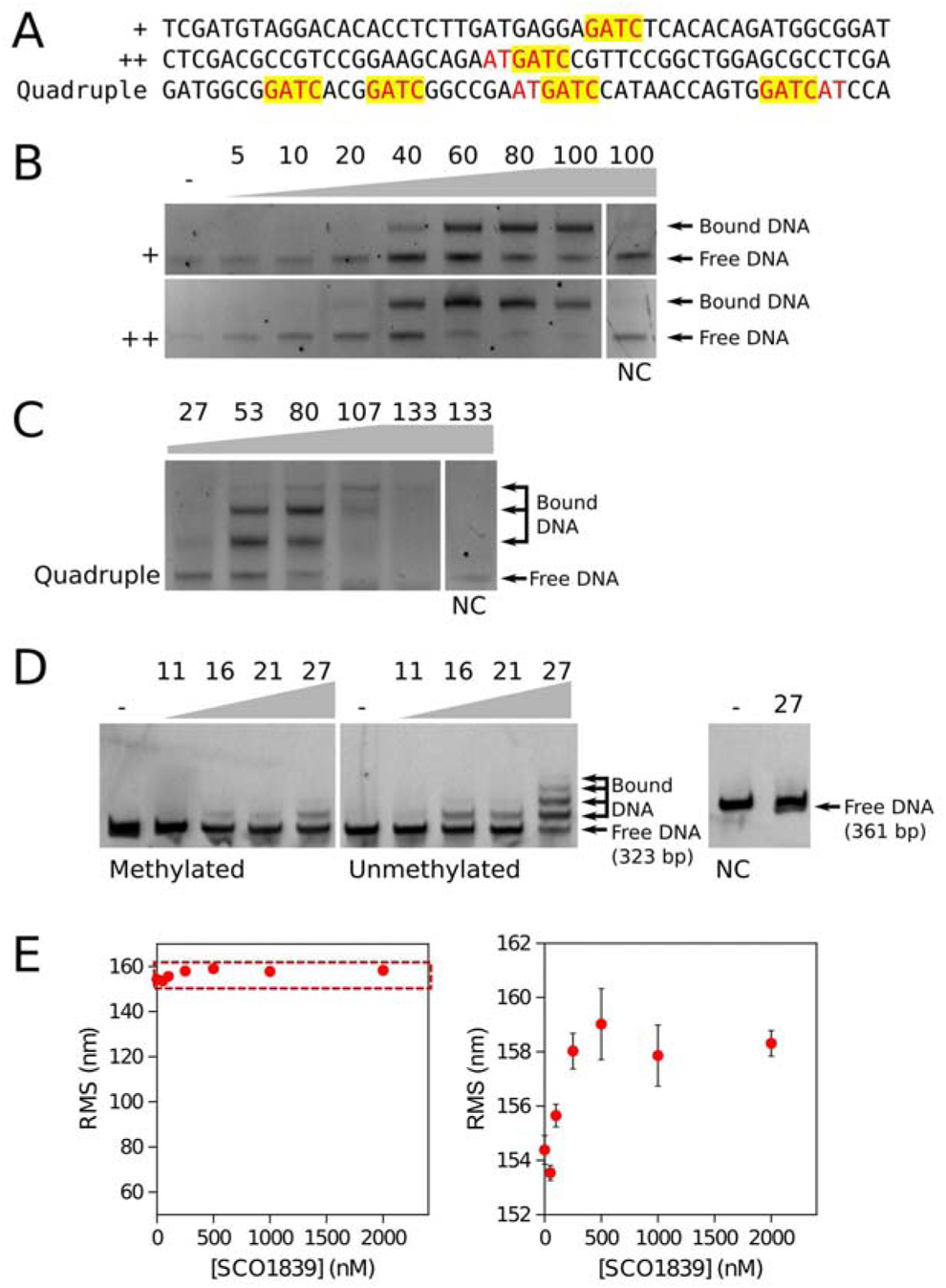
Binding specificity and affinity of Gbn protein. **(A)** Sequence of short DNA fragments used in the Electrophoretic mobility shift assays (EMSA). The motifs included in the EMSA sequences were designed with different degrees of binding strength, GATC motif is indicated by “+”, GATCWT by “++”, and the DNA sequence containing two GATCWTs and two GATCs is indicated by “Quadruple”. GATC motifs are highlighted with the trailing (A/T)T sequence coloured red. **(B, C)** Electrophoretic mobility shift assays (EMSA) using 6×His-tagged Gbn on synthetic 50 bp dsDNAs containing the different binding strengths GATC motifs described in (A). **(D)** EMSA using 6×His-tagged Gbn protein performed on DNA extracted from DNA methylation-deficient *E. coli* ET12567 and methylation-positive *E. coli* JM109. The *gbn* promoter region (−561 to −239) contains eight GATC motifs and was used as test sequence. Negative Control (NC) sequence derived from a random location with no detectable Gbn protein binding as shown by ChIP-Seq analysis, specifically the position 5,052,200 to 5,052,548 (349 bp). In panels B), C), and D), the protein-to-DNA molar ratios are shown on top of each gel picture, 10 μL of mixed samples was loaded per lane with DNA concentrations: (B) 5 nM; (C) 7.5 nM; and (D) 2 nM. **(E)** Root mean square displacement (RMS) of the bead for Gbn on a 685 bp DNA substrate containing the *gbn* promoter region as shown as a function of protein concentration. Error bars indicate the standard error of the data points (N ≥ 100). On the right, a close-up of the dashed region is shown.

The GATC motif is the target sequence for deoxyadenosine methylase (DAM), which is essential for DNA mismatch repair in E. coli (62). Another DNA methylation takes place on the second deoxycytosine of the sequence CCTGG (DCM) (63). *S. coelicolor* lacks both methylation systems and degrades methylated exogenous DNA (51, 64). Which proteins are involved in the recognition and restriction system in Streptomyces remains unknown (65, 66). To test whether Gbn play a role in this restriction system, we compared the transformation efficiency of methylated DNA and non-methylated DNA in the *gbn* null and in the parental strain M145. No significant differences were observed in the transformation efficiencies between parent and *gbn* mutant, which suggests that Gbn does not play a role in restriction of methylated DNA (data not illustrated). Next, we tested the affinity difference of Gbn for GATC and GA^m^TC in an electrophoretic mobility shift assay (EMSA). The results revealed that Gbn had lower affinity for methylated DNA (Figure 5D), which may be caused by a steric effect of the methyl group on Gbn binding.

### Gbn does not alter the conformation of the DNA

In order to investigate whether Gbn alters DNA conformation in vitro, we performed Tethered Particle Motion (TPM) experiments (67). If the binding of a protein to DNA induces bends or affects DNA stiffness this translates into a reduction, or an increase of the root mean square displacement (RMS) respectively compared to that of bare DNA (68–70). Here we used a 685 bp DNA substrate containing the *gbn* promoter region with 10 GATC(WT) sites. Addition of the Gbn protein resulted in a very small increase in RMS of around 4 nm at ≥ 250 nM (Figure 5E). This could be explained by Gbn occupying the 10 binding sites without causing major changes in the DNA conformation. While Gbn does not deform DNA promoting compaction, the increase in RMS is indicative of the DNA being somewhat stiffened by binding of the protein. Earlier studies indicate that the observed mild increase in RMS corresponds to an increase in persistence length of about 5 nm as compared to bare DNA (70). Note that qualitatively similar effects have been observed for E. coli and P. aeruginosa H-NS like proteins (68, 69). These properties suggest that rather than contributing directly to organization and compaction of the chromosome, Gbn functions in regulation of genome transactions such as transcription.

### Gbn binding events are found in the regulatory regions of some 10% of all genes

To obtain better insight into which genes are affected by the binding of Gbn, we investigated Gbn binding events in the putative promoter regions (−350 to +50 relative transcription start site of all genes). In total, Gbn bound to the promoter of 769 genes at both 25 h and 48 h (Table S1). Of these genes, 44.5% (342 genes) had at least binding event with more than 10-fold enrichment in the corresponding promoter region. These genes include many genes from cell division and cell wall (dcw) cluster (SCO2077-SCO2088), 50S ribosomal protein gene cluster (SCO4702-SCO4727), tRNAs (SCOt02-SCOt50), atrA, redY, SCO1311 and *gbn* itself (Table 2, Figure 4B). One Gbn binding event was found upstream of ssgR, with a peak at around nt position −483 relative to the translation start site (Figure 4C). This is in accordance with the previous observation that Gbn binds to the ssgR promoter region (22). Gbn not only bound to the promoter regions of five tRNA genes, but also to the promoter region of SCO1311, which has a tRNA-editing domain responsible for hydrolysing mis-acylated tRNA (coverage 68%, Pfam: PF04073). Interestingly, the promoter regions of SCO1311 and SCOt32 showed the largest difference in Gbn binding between 25 h and 48 h samples, with three times higher enrichment at 25 h than at 48 h (Figure 4D, F, Table 2). SCOt23 specifies a leucyl-tRNA with anticodon UAG, which is required for the translation of the rare leucine codon CUA. The rarest codon in *S. coelicolor* is another leucine codon, namely UUA. The tRNA recognizing the UUA codon is specified by bldA, and the corresponding TTA codon occurs specifically in many genes involved in development and antibiotic production, making those genes bldA dependent (71–74). The CTA leucine codon is also very rare in Streptomyces genes, representing only 0.31% of all leucine codons in *S. coelicolor*. In this light, it would be interesting to see if SCOt23 also plays a role in the control of developmental gene expression, and what the role is of Gbn in the control of SCOt23 transcription.

**Table 2.** Genes with strong Gbn binding event at the promoter region.

| Gene ID | Gene product | Binding width |  | Binding summit position** |  | Fold enrichment |  |
| --- | --- | --- | --- | --- | --- | --- | --- |
|  |  | 25h | 48h | 25h | 48h | 25h | 48h |
| SCO1311* | hypothetical protein with tRNA edit domain | 271 | ND<sup>†</sup> | -76 | -72<sup>†</sup> | 13.41 | 2.42<sup>†</sup> |
| SCO1839* | transcriptional regulator | 547 | 530 | -326 | -324 | 67.75 | 58.18 |
| SCO2077* | DivIVA | 251 | 300 | -90 | -91 | 17.19 | 13.48 |
| SCO2081* | YlmD | 271 | 378 | -348 | -348 | 10.44 | 17.18 |
| SCO2084* | MurG (UDPdiphospho-muramoylpentapeptide beta-N-acetylglucosaminyltransferase) | 280 | 313 | -280 | -273 | 13.22 | 13.41 |
| SCO2086* | MurD (UDP-N-acetylmuramoyl-L-alanyl-D-glutamate synthetase) | 482 | 471 | -90 | -100 | 9.63 | 10.62 |
| SCO2088* | MurF (UDP-N-acetylmuramoylalanyl-D-glutamyl-2, 6-diaminopimelate- D-alanyl-alanyl ligase) | 227 | 285 | -5 | -10 | 7.37 | 14.21 |
| SCO3213 | TrpG (anthranilate synthase component II) | 247 | 299 | -240 | -236 | 7.85 | 10.69 |
| SCO3224 | ABC transporter ATP-binding protein | 349 | 367 | +33 | +48 | 14.23 | 15.90 |
| SCO3225 | AbsA1 (two component sensor kinase) | Δ | Δ | -168 | -183 | Δ | Δ |
| SCO4702 | RplC (50S ribosomal protein L3) | 459 | 511 | -34 | -37 | 12.19 | 22.16 |
| SCO4707 | RplV (50S ribosomal protein L22) | 290 | 388 | -198 | -196 | 15.80 | 25.17 |
| SCO4713 | RplX (50S ribosomal protein L24) | 372 | 406 | -7 | -5 | 58.66 | 44.69 |
| SCO4714 | RplE (50S ribosomal protein L5) | Δ | Δ | -330 | -328 | Δ | Δ |
| SCO4717 | RplF (50S ribosomal protein L6) | 372 | 418 | -324 | -326 | 25.78 | 23.66 |
| SCO4718 | RplR (50S ribosomal protein L18) | 252 | 641 | +46 | +45 | 10.27 | 19.35 |
| SCO4721 | RplO (50S ribosomal protein L15) | 380 | 432 | +22 | +21 | 34.82 | 29.13 |
| SCO4726 | RpmJ (50S ribosomal protein L36) | 310 | 761 | -3 | +21 | 6.69 | 15.32 |
| SCO4727 | RpsM (30S ribosomal protein S13) | Δ | Δ | -305 | -282 | Δ | Δ |
| SCO5880* | RedY (RedY protein) | 286 | 347 | -168 | -161 | 14.59 | 18.85 |
| SCOt02 | tRNA Val (anticodon CAC) | 509 | 559 | -136 | -138 | 56.47 | 55.65 |
| SCOt17 | tRNA Gly (anticodon UCC) | 295 | 348 | -325 | -320 | 20.40 | 28.92 |
| SCOt23* | tRNA Leu (anticodon UAG) | 338 | 280 | -78 | -90 | 25.89 | 8.10 |
| SCOt49 | tRNA Thr (anticodon GGU) | 261 | 294 | -170 | -168 | 15.21 | 13.95 |
| SCOt50 | tRNA Met (anticodon CAU) | Δ | Δ | -289 | -287 | Δ | Δ |

\* Shown in Figure 4
\*\* Relative to the start of the gene (+1)
<sup>†</sup> Binding region not detected by the software, local summit position and corresponding fold enrichment are shown instead
Δ same as above, shared binding region as the gene above

Interestingly, many of the strongest enrichments of binding in promoter regions (15- to 20-fold) were found in the dcw cluster, which encompasses many key genes for cell division and cell-wall synthesis, including genes for the cell division scaffold FtsZ and DivIVA which is essential for growth (Figure 4G). However, transcriptomics data shows that the expression of dcw gene cluster was not significantly altered by deletion of *gbn*. This may be explained by the fact that NAPs typically affect global gene expression through remodelling of DNA structure and not by direct activation or repression of transcription (21). Interestingly, strong binding was also observed in ribosomal protein operons (Figure 4H). In Streptomyces, development and the regulation of secondary metabolism are associated with changes in the expression of ribosomal proteins (75, 76). This phenomenon may be related with the resistance mechanism of many antibiotics (77, 78).

### Deletion of *gbn* accelerates sporulation of *S. coelicolor*

To obtain insights into the possible role of Gbn in the life cycle of *S. coelicolor*, a knock-out mutant was generated using a strategy published previously (79). For this, the +1 to +207 region of the gene was replaced by the apramycin resistance cassette aac(C)IV, which was flanked by loxP sites. The loxP sites allowed removal of the cassette using Cre recombinase, important to minimise polar effects. To genetically complement the mutant and see if the wild-type phenotype would be restored, the −565/+228 region of *gbn* was amplified from the *S. coelicolor* chromosome and cloned into pHJL401, a shuttle vector that is useful for genetic complementation due to its low copy number in streptomycetes (80). To also analyse the effect of enhanced expression of *gbn*, a second strain was constructed using CRISPR-Cas9 wherein the native promoter of *gbn* (−157 to +4, start codon modified to ATG) was replaced by the strong constitutive ermE promoter (81). For details see the Materials and Methods section.

The morphology of *gbn* null mutant did not show significant changes comparing to wild type, except for reduced production of the blue-pigmented antibiotic actinorhodin (Figure S1). However, the mutant showed slightly accelerated development in comparison to the parental strain. To investigate this altered timing of development in more detail, time-lapse imaging was performed on confluent mycelial lawns. This method allows monitoring multiple morphological characteristics and in particular quantifying differences in timing of the individual developmental stages, based on pigmentation of the mycelia. When aerial hyphae are formed, the brightness increases due to the increased density of colourless hyphae. The brightness decreases when grey-pigmented spores are produced (see Material and Methods section for details). Deletion of *gbn* led to a 2-5 h acceleration of development as compared to the parental strain (Figure 2A). 54 h after inoculation, the light intensity of the *gbn* mutant again increased (Figure 2A), which may be due to premature germination and renewed growth. The acceleration of development was reversed when a wild-type copy of *gbn* was re-introduced into the null mutant, a genetic complementation experiment that confirmed the accelerated development was indeed primarily due to the deletion of *gbn*. Conversely, the enhanced and constitutive expression of *gbn* delayed sporulation by approximately 17 h (Figure 2A). Taken together, this strongly suggests that the expression of *gbn* correlates to the timing of sporulation.

**Figure 2.**
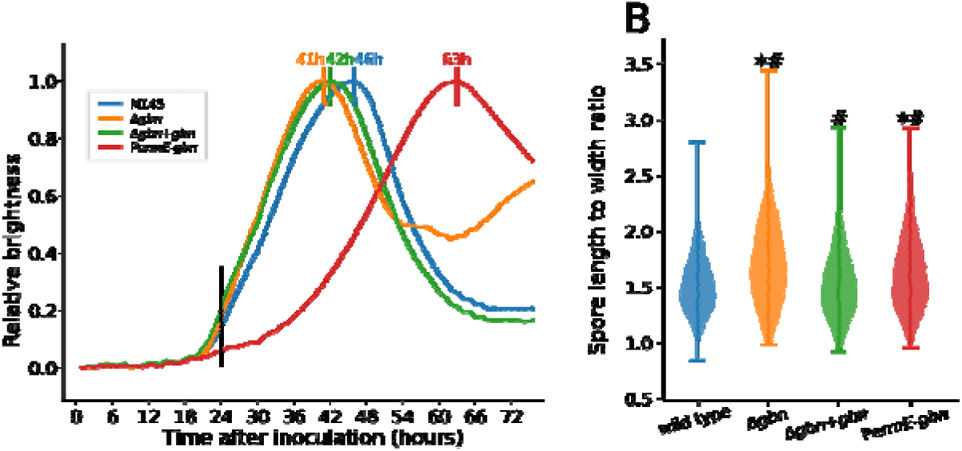
Growth and spore shape analysis of *gbn* mutants. **(A)** scanner measurements of confluent plate brightness of *S. coelicolor* M145 and mutants (Δ*gbn*, Δ*gbn*+*gbn*, P*ermE*-*gbn*) grown as confluent lawns. The X-axis represents the time after inoculation and the Y-axis the normalized brightness measured from time lapse scan pictures. Mutants are presented in different colours; the timing of curve peaks are marked on top. Black vertical line illustrates 24 h time point. **(B)** Violin plot showing the distributions of the spore length to width ratio as derived from SEM images. Each strain was grown on MM agar for 5 days before imaging. with significantly (p < 0.01) different length to width ratio compared to *S. coelicolor* M145 (Mann–Whitney U test), # with significantly (p < 0.01) different variance compared to *S. coelicolor* M145 (Levene’s test).

Closer examination of the spores by scanning electron microscopy (SEM) revealed an increased length-to-width ratio of the spores in both the deletion mutant and the strain with enhanced expression of *gbn* (Mann-Whitney U test, p-value < 0.01, Table 1, Figure 2B). The complemented strain produced spores with a normal ratio. The statistical variation in the spore length-to-width ratio showed a larger variance in the engineered strains as compared to that of the parental strain (Levene’s test, p-value < 0.01), with the *gbn* deletion mutant showing the largest variance. Complementation of the mutant reduced the variance to a level close to that of the wild type. Thus, deletion of *gbn* altered both the timing of sporulation and the morphology of the spores.

**Table 1.** Spore length to width ratio of all strains comparing with parent strain M145

| Strain | n | Median | Mann–Whitney *U* test of difference |  | Variance | Levene's test of equal variance |  |
| --- | --- | --- | --- | --- | --- | --- | --- |
|  |  |  | *U* | *p*-value |  | *W* | *p*-value |
| M145 | 560 | 1.49 | - | - | 0.069 | - | - |
| *Δgbn* | 332 | 1.69 | $1.2 \times 10^5$ | $9.3 \times 10^{-16}$ | 0.145 | 49.51 | $3.93 \times 10^{-12}$ |
| *Δgbn+gbn* | 556 | 1.50 | $1.6 \times 10^5$ | 0.40 | 0.099 | 13.49 | $2.52 \times 10^{-4}$ |
| PermE-*gbn* | 441 | 1.60 | $1.5 \times 10^5$ | $1.4 \times 10^{-7}$ | 0.125 | 34.89 | $4.78 \times 10^{-9}$ |

### Correlation between Gbn binding and gene expression

To obtain insights into the effect of Gbn on global gene expression, we performed transcriptomics analysis, comparing transcriptional changes between the *gbn* mutant with its parent during vegetative and aerial growth. For this, strains were grown on minimum agar medium covered with cellophane and RNA samples were prepared in triplicate after 24 h (vegetative growth) and 45 h (aerial growth). A table with all read counts for each gene can be found at GEO accession GSE186136; differential expression analysis table can be found in Table S2.

Table 2 lists the expression of all genes associated with strong Gbn binding in their promoter regions. This surprisingly revealed no significant changes between parent and *gbn* mutant, indicating that Gbn does not directly affect the level of transcription of target genes. As an example, Gbn binds to the promoter region of atrA (Figure 4E), a global regulatory gene which among others trans-activates the transcription of actII-ORF4, the cluster-situated activator of the actinorhodin biosynthetic gene cluster (82). Binding may be reciprocal, as an AtrA binding site was identified in the *gbn* promoter region using the PREDetector algorithm (83), with a high confidence score of 13.3, similar to that for the AtrA binding element upstream of actII-ORF4. However, atrA transcription did not change in the *gbn* null mutant. In total 13 binding regions were detected in the red cluster at both time points, with a very strong binding site upstream of redY (Figure 4H). Again, transcription was not altered significantly between wild-type and *gbn* mutant.

Interestingly, the average level of gene expression during vegetative growth was significantly higher than during aerial growth for genes with Gbn binding in promoter regions, which was even more pronounced in a *gbn* mutant background (Table 3). Such an effect was not seen for genes where Gbn bound only to the coding regions (+50 to end of gene). Thus, binding of Gbn to the promoter regions of genes has a subtle but significant suppressive effect on overall gene expression, which cannot be explained by direct effect on the expression of specific genes.

**Table 3.**
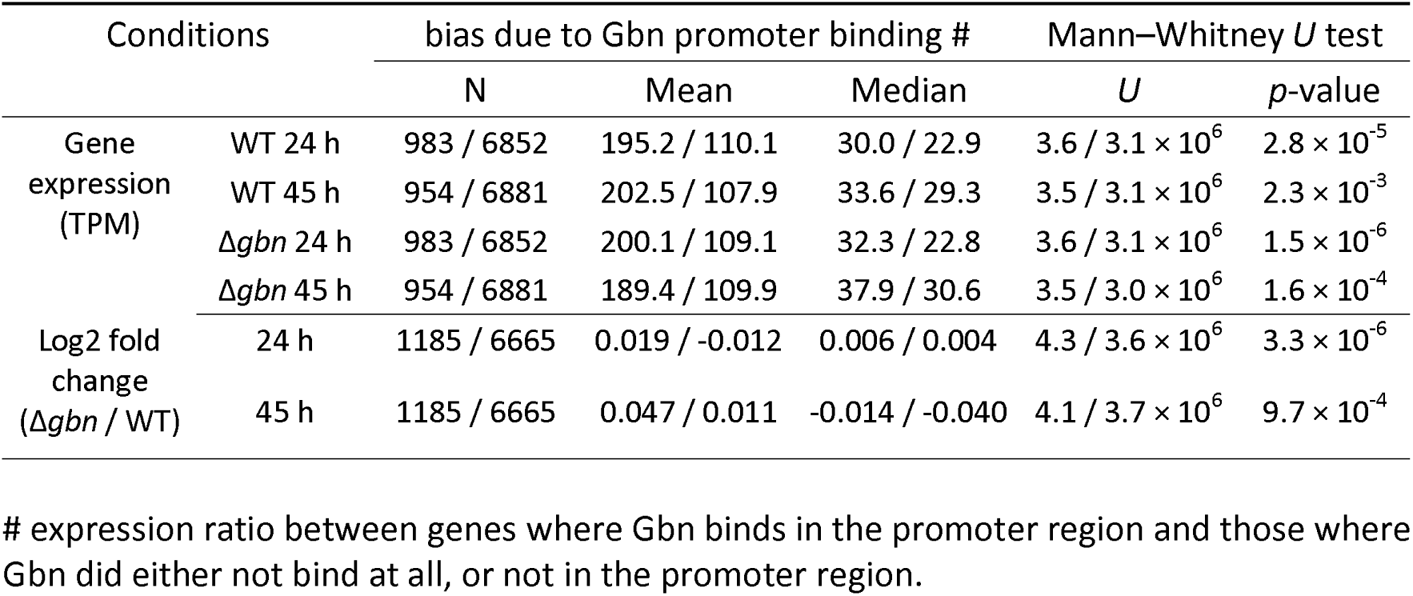
Expression differences between genes where Gbn bound in the promoter region.

| Conditions |  | bias due to Gbn promoter binding # |  |  | Mann–Whitney *U* test |  |
| --- | --- | --- | --- | --- | --- | --- |
|  |  | N | Mean | Median | *U* | *p*-value |
| Gene expression (TPM) | WT 24 h | 983 / 6852 | 195.2 / 110.1 | 30.0 / 22.9 | $3.6 / 3.1 \times 10^6$ | $2.8 \times 10^{-5}$ |
| | WT 45 h | 954 / 6881 | 202.5 / 107.9 | 33.6 / 29.3 | $3.5 / 3.1 \times 10^6$ | $2.3 \times 10^{-3}$ |
| | $\Delta gbn$ 24 h | 983 / 6852 | 200.1 / 109.1 | 32.3 / 22.8 | $3.6 / 3.1 \times 10^6$ | $1.5 \times 10^{-6}$ |
| | $\Delta gbn$ 45 h | 954 / 6881 | 189.4 / 109.9 | 37.9 / 30.6 | $3.5 / 3.0 \times 10^6$ | $1.6 \times 10^{-4}$ |
| Log2 fold change ($\Delta gbn$ / WT) | 24 h | 1185 / 6665 | 0.019 / -0.012 | 0.006 / 0.004 | $4.3 / 3.6 \times 10^6$ | $3.3 \times 10^{-6}$ |
| | 45 h | 1185 / 6665 | 0.047 / 0.011 | -0.014 / -0.040 | $4.1 / 3.7 \times 10^6$ | $9.7 \times 10^{-4}$ |

Transcriptomics data showed a strong increase in the transcription of *gbn* itself at 45 h as compared to 24 h (Figure 6A). This increased expression of *gbn* during development was also seen in transcriptome data published by others; all public datasets showed an increase in *gbn* expression over time and in a medium-independent manner (14, 75, 84, 85). Interestingly, repeated ChIP-seq experiments on 24 h old samples or younger failed to pull down any DNA, while after 25 h, some DNA was immunoprecipitated. This correlates to the observation that Gbn expression is initiated after around 24 h after start of growth.

**Figure 6.**
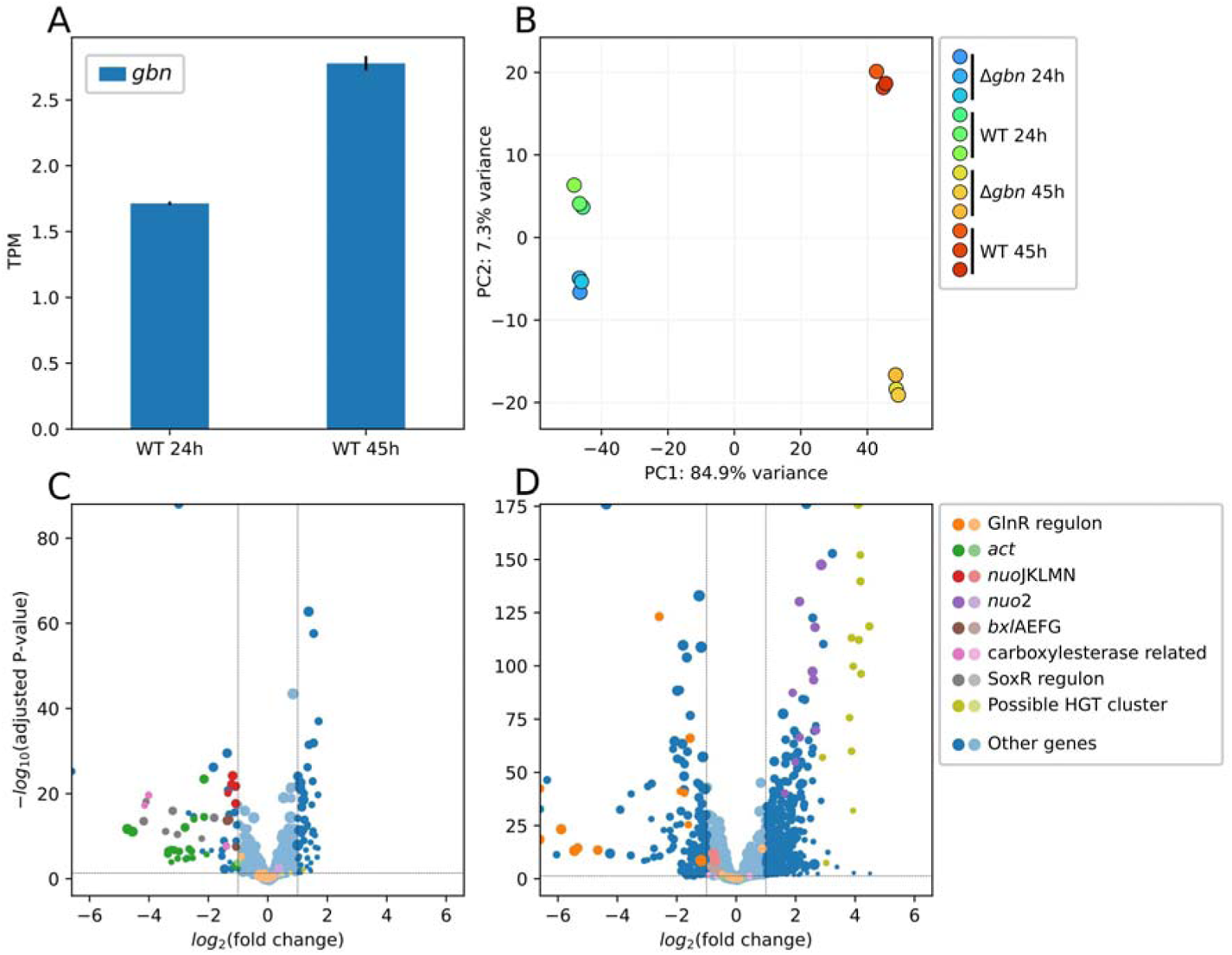
Transcriptomics data of vegetative (24 h) and aerial (45 h) growth stage cells of Δ*gbn* and wild type strain. **(A)** Expression of *gbn* gene in wild type strain. Error bars indicate the standard error of triplicates. **(B)** Principal component analysis of transcriptomics samples with *gbn* excluded from analysis. **(C, D)** Volcano plot showing changes in gene expression due to deletion of *gbn* (Δ*gbn* vs. wild type) at vegetative (C) and aerial growth stage (D). Interested genes are highlighted in colours other than blue. Both (C) and (D) share the same colour scheme which is shown in the legend to the right. Lighter colours indicating the change is not significant (fold change ≤ 2, p-value ≥ 0.05); size of each dot represents the normalized average expression of related conditions. Except (A), all plot data was calculated using DESeq2 with shrinkage function (55, 56).

### Gbn stimulates secondary metabolic pathways

Principal component analysis (PCA) of transcriptomics data showed an increased distance for aerial growth stage samples (Figure 6B). This suggests that deletion of *gbn* leads to more fundamental transcriptomic changes at a later growth stage. In the *gbn* null mutant, many secondary metabolomic genes were down-regulated, especially during vegetative growth phase (Table 4, Figure 6C), and most of these genes did not show Gbn binding. The down-regulated genes include those of the act BGC (SCO5071-SCO5092) and of the SoxR regulon, which can be activated by γ-actinorhodin (86). This is consistent with the observed slower blue pigmentation of the mutant on SFM agar plates (Figure S1). Additionally, genes for a carboxylesterase regulon (SCO0319-SCO0321) and a xylobiose transporter regulon (bxlEFGA; SCO7028-SCO7031) (87) were also down-regulated.

**Table 4.**
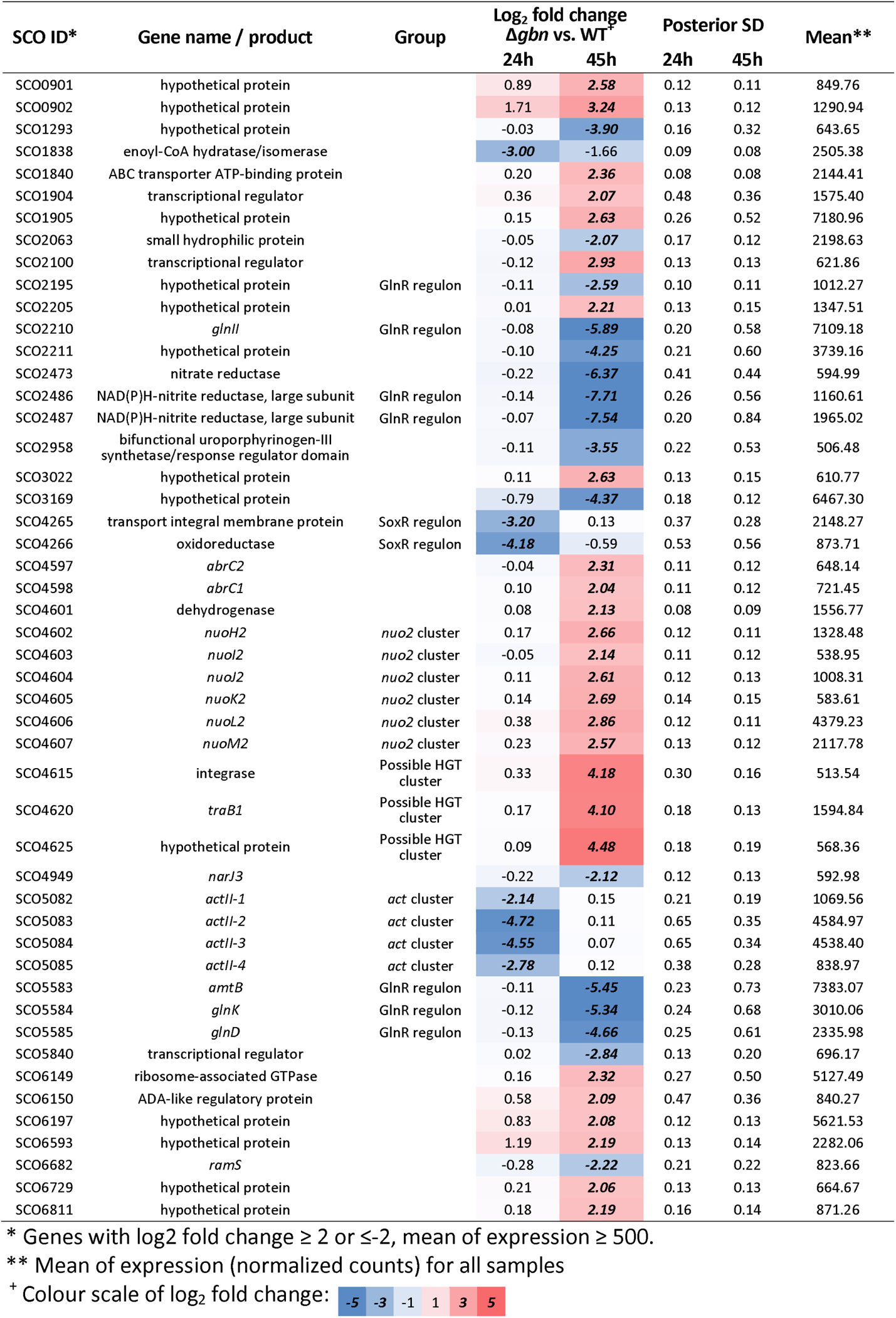
Genes with high expression that changes the most by deletion of *gbn*

During aerial growth (45 h), many more genes are affected in the *gbn* null mutant (Table 4, Figure 6D), which correlates well with the increased PCA distance at this time point. An obvious category of down-regulated genes belongs to the GlnR regulon (gene list obtained from (88)), especially for SCO2195, glnII (SCO2210), nirB (SCO2486, SCO2487), amtB (SCO5583), glnKD (SCO5584, SCO5585) (Table 4). Among the up-regulated genes during aerial growth in the mutant, SCO4615-SCO4627 were most strongly up regulated. These include phage-related genes, encoding among others an integrase (SCO4615), an excisionase (SCO4616), and the DNA transfer-related TraA1 (SCO4621), TraB1 (SCO4620) and SpdB2 (SCO4625) (89). In the vicinity of this gene cluster lie more genes that are significantly up regulated in the *gbn* mutant during aerial growth. These genes include an atypical two-component system that supresses antibiotic production (SCO4596-SCO4598) (90) and the second NADH dehydrogenase I gene cluster (nuo2; SCO4599-SCO4608). The nuo2 cluster lacks genes for NuoCDEFG and L subunits and its function is unclear. Conversely, nuoJKLMN (SCO4571-SCO4575) from the canonical nuo gene cluster (91) were down-regulated in the *gbn* mutant (Table 4, Figure 6D). The gene products of the nuo2 cluster may cooperates with the proteins derived from the canonical nuo operon. The altered balance in the expression of components of the NADH dehydrogenase complex derived from the nuo and nuo2 operons may lead to a hybrid machinery. Whether this is indeed the case, and how this affects energy production during development, remains to be elucidated.

In conclusion, transcriptomics data confirmed the early sporulation and suppressed secondary metabolic processes in *gbn* null mutant. It is tempting to propose that these two processes have causal relationship that directly related with Gbn. However, direct links needs to be found to confirm this relationship and understand the regulation pathway that drives these changes related to deletion of *gbn*.

### Summary

Gbn (SCO1839) protein is a conserved NAP among Actinobacteria species, it is a highly pleiotropic DNA binding protein that plays a role in the (timing of) development and antibiotic production of *S. coelicolor*. Strains in which *gbn* had been deleted or over-expressed showed accelerated and delayed sporulation, respectively, suggesting that Gbn plays a role in the accurate timing of development. Both in vivo and in vitro experiments revealed that Gbn binds specifically to a GATC DNA motif and especially those followed by either AT or TT. In addition, we have shown that the methylation of adenine in the GATC sequence reduced affinity of Gbn for its binding sites. Transcriptomics analysis showed Gbn has a suppressive effect on the genes that bound by Gbn on their promoter regions. This suppressive effect, together with thousands genome locations that Gbn binds to, may instead leads to stimulation of secondary metabolic pathways and postponed sporulation.

These results described Gbn as a representative member of a new NAP family that might play important roles in the development and antibiotics production in streptomycetes.

## DATA AND CODE AVAILABILITY

Clean ChIP-Seq reads and binding region identification (peak calling) files are available at GEO database (92) with accession number GSE165795. [For reviewers: go to https://www.ncbi.nlm.nih.gov/geo/query/acc.cgi?acc=GSE165795, enter token ujulcouyzxcfhmn into the box]. Clean RNA-Seq reads and gene read-counts tables are available at GEO database with (92) with accession number GSE186136. [For reviewers: go to https://www.ncbi.nlm.nih.gov/geo/query/acc.cgi?acc=GSE186136, enter token qryzegmcrxifrqf into the box]. Complete ChIP-Seq analysis and transcriptomics analysis code and related data table used in this research can be found at https://github.com/snail123815/Gbn-the-SNP-publication-scripts. Time lapse scanner image analysis code can be found at https://github.com/snail123815/scanLapsePlot.

## ACKNOWLEDGMENTS

We are grateful to Kenny McDowall for stimulating discussions.

