## Supplemental figures and tables for "Systems-wide analysis of the GATC-binding nucleoid-associated protein Gbn and its impact on *Streptomyces* development"

belonging to the manuscript

**Table S1.** Gbn binding regions that overlap with gene promoter regions
Complete table can be accessed: Supplementary Table S1.xlsx.

**Table S2.** Transcriptomics differential expression analysis result
Complete table can be accessed: Supplementary Table S2.xlsx.

**Table S3.** Bacterial strains used in this study

| Strain | Genotype/description | Reference/vendor reference |
|---|---|---|
| *E. coli* JM109 | See reference | (37) |
| *E. coli* ET12567 | See reference | (27) |
| *E. coli* ET12567/pUZ8002 | See reference | (62) |
| *E. coli* BL21 CodonPlus (DE3)-RIPL | See reference | Agilent 230280 |
| *S. coelicolor* A3(2) M145 | See reference | (29) |
| GAD003 | M145Δ*gbn* | This study |
| GAD014 | M145Δ*gbn* + pGWS1260 | This study |
| GAD039 | Part of *gbn* promoter replaced by PermE | This study |
| GAD043 | 3×FLAG fused to *gbn* 3’-term | This study |

**Table S4.** Primers used in this study

| Purpose | No. | Name | Sequence 5'-3' | Note |
|---|---|---|---|---|
| SCO1839 knock-out and complementation | 1 | SCO1839FU_F | CGATGGATCCGACGCCGACTCGATCATCTG | |
| | 2 | SCO1839FU_R | CGATTCTAGACCGTGCCTCCTCATGGGAAG | |
| | 3 | SCO1839FD_F | CGATTCTAGAGCCACTTCGGCCTGACGTCC | |
| | 4 | SCO1839FD_R | CGATAAGCTTCCGGCCTGCTCGACGAAAC | |
| | 5 | SCO1839CM_F | ACTGAAGCTTGGGACGTCAGGCCGAAGTG | |
| | 6 | SCO1839CM_R | ACTGGGATCCGGTCTCCTCGGCCAACGTG | |
| SCO1839 over-expression | 7 | 1839_Ukp_F | CCGATCTAGACGACGCCGACTCGATCATCTG | No.1 changed to XbaI |
| | 8 | 1839_Ukp_R | CAGTGAGCTCTCCGCCGAACGAGTTTCTCC | |
| | 9 | SCO1839OP_F | CAGTCATATGGCCGAGACTCTGAAGAAGGG | |
| | 10 | SCO1839FD_R_XbaI | CGATTCTAGACCGGCCTGCTCGACGAAAC | No.4 changed to XbaI |
| | 11 | Sp_1839U_F | ACGCTCGGCAAGGCTGATGAGACA | Spacer assembly |
| | 12 | Sp_1839U_R | AAACTGTCTCATCAGCCTTGCCGA | |
| 3×FLAG tag knock-in | 13 | 1839sp_down_F | ACGCGCCACTTCGGCCTGACGTCC | Spacer assembly |
| | 14 | 1839sp_down_R | AAACGGACGTCAGGCCGAAGTGGC | |
| | 15 | 1839flagFL_UR | AGTCGCCGTCGTGGTCCTTGTAGTCGGCCGAAGTGGCCTTCTTGC | |
| | 16 | 3xFLAG+1839ending | GACTACAAGGACCACGACGGCGACTACAAGGACCACGACATCGACTACAAGGACGACGACGACAAGTGA CGTCCCGGGCGGCCCCCGGCTCCGAAAGTGACGGTGGCCACCCGGT | |
| | 17 | 1839flagCR_DR | GCGGCCTTTTTACGGTTCCTGGCCTCCTCGTAGTCCCGCGTCTGG | |
| | 18 | 1839flagCR_UF_xbaI | CGATTCTAGATGGTCTCCTCGGCCAACGTG | |
| | 19 | 1839flagCR_DR_xbaI | CGATTCTAGACCTCGTAGTCCCGCGTCTGG | |
| 50 bp EMSA fragments | 24 | p1839_50Single_strong | CTCGACGCCGTCCGGAAGCAGAATGATCCGTTCCGGCTGGAGCGCCTCGA | |
| | 25 | p1839_50Single_strong_R | TCGAGGCGCTCCAGCCGGAACGGATCATTCTGCTTCCGGACGGCGTCGAG | |
| | 26 | p1839_50Single_weak | TCGATGTAGGACACACCTCTTGATGAGGAGATCTCACACAGATGGCGGAT | |
| | 27 | p1839_50Single_weak_R | ATCCGCCATCTGTGTGAGATCTCCTCATCAAGAGGTGTGTCCTACATCGA | |

| | | | | |
|---|---|---|---|---|
| 50 bp EMSA fragments | 28 | p1839_50Quadruple | GATGGCGGATCACGGATCGGCCGAATGATCCATAACCAGTGGATCATCCA | |
| | 29 | p1839_50Quadruple_R | TGGATGATCCACTGGTTATGGATCATTCGGCCGATCCGTGATCCGCCATC | |
| | 30 | npio_B50 | GCAATTCGTAGGACGGAAGTGCGCAGGGTGCCTCAAATGCGGCCCTATAG | Negative control |
| | 31 | npio_B50_R | CTATAGGGCCGCATTTGAGGCACCCTGCGCACTTCCGTCCTACGAATTGC | |
| Long EMSA fragments for possible methylation affection | 32 | p1839_ip_F | CCGCCGAACGAGTTTCTCC | *Eco*RI site inside PCR product, the fragment was digested using *Eco*RI and HindIII for inserting into pUC19 |
| | 33 | p1839_ip_R_HindIII | GCATAAGCTTCCTCGGCCAACGTGCTG | |
| | 34 | noip_A_F_EcoRI | GCATGAATTCGCCGCCGTTCGGTGGTGTC | Negative control |
| | 35 | noip_A_R_HindIII | GCATAAGCTTGCCAGCGAGGCCCGCTTC | |
| TPM | 36 | puc19_TPM2_F | ACCTATTCCGTTTCAGCCAGCAGACAAGCTGTGACCGTCT | |
| | 37 | puc19_TPM2_R | CCGGTTCTCGCTGAGCTCCTGGGGTGCCTAATGAGTG | |
| | 38 | TPM2_p1839_3_F | CTGGCTGAAACGGAATAGGTGTAACCCGGCTGCCCTTCTTCA | |
| | 39 | TPM2_p1839_3_R | AGCTCAGCGAGAACCGGACAGCTCTCCGGCGGCGAGAAGAC | |
| | 40 | TPM_F_biotin | Biotin-CTGGCTGAAACGGAATAGGT | |
| | 41 | TPM_R_digoxygenin | Digoxygenin-AGCTCAGCGAGAACCGG | |
| Promoter probing | 42 | pSCO1839_F_BamHI | CGATGGATCCCTCATGGGAAGTGCGCTCTG | |
| | 43 | pSCO1839_R_SacI | ACGTGAGCTCTCCGTGATCCGCCATCTGTG | |
| qPCR | 44 | SCO1839_QF | GAGCATGCGGTGCACAAAG | |
| | 45 | SCO1839_QR | GCGGCAGACCTGAAGAAGAAG | |
| | 46 | SCO3873_qF | GCGTGGTGGACACGAAGAAG | Inner control |
| | 47 | SCO3873_qR | GCACCAAGACCGACGACTAC | |
| | 48 | SCO5359_qF | TACGTCGAGACGCAGGTCAG | Inner control |
| | 49 | SCO5359_qR | CTGCTTGCCCGTGTAGAAC | |

Underlined characters indicating restriction sites or overhangs for assembly.
GGATCC, *Bam*HI; GAATTC, *Eco*RI; AGGCCT, *Stu*I; TCTAGA, *Xba*I; GAGCTC, *Sac*I; CATATG, *Nde*I; AAGCTT, *Hind*III

**Table S5.** Plasmids and constructs used in this study

| Plasmid and construct | Description | reference/vendor ID |
|---|---|---|
| pWHM3 | *E. coli*/*Streptomyces* shuttle vector, high copy number and unstable in *Streptomyces* | (31) |
| pUWL-Cre | *E. coli*/*Streptomyces* shuttle vector expressing the Cre recombinase in *Streptomyces* | (33) |
| pHJL401 | *E. coli*/*Streptomyces* shuttle vector, 5-10 copies per chromosome in *Streptomyces* | (80) |
| pHM10a | *E. coli*/*Streptomyces* shuttle vector, designed for gene over-expression using consecutive promoter P*ermE* | (36) |
| pCRISPomyces-2 | *E. coli*/*Streptomyces* shuttle vector, harbouring codon optimised *cas9*, designed for easy inserting spacer sequences. Recombination template is designed to be inserted at *Xba*I site. | (35) |
| pCRISPR-Cas9 | *E. coli*/*Streptomyces* shuttle vector, harbouring codon optimised *cas9*, designed for easy inserting spacer sequences. Recombination template is designed to be inserted through in-vitro assembly to *Stu*I site | (81) |
| pGWS728 | Construct harbouring *aac(3)IV* | (32) |
| pET28a | *E. coli* vector, designed to build His-tag fusion protein expression construct | Novagene 69864-3 |
| pUC19 | *E. coli* vector with multi-copy origin of replication | NEB N3041 |
| pGWS1255 | pWHM3 containing flanking regions of *gbn* with apramycin resistance cassette with *loxP* sites inserted as *Xba*I fragment between flanking regions | This study |
| pGWS1260 | pHJL401 harbouring *gbn* and its own promoter region | This study |
| pGWS1298 | pCRISPomyces-2 with spacer sequence from near the end of *gbn*, containing recombination template for 3×FLAG tag knock-in | This study |
| pGWS1295 | pCRISPomyces-2 with spacer sequence from near the beginning of *gbn*, containing recombination template for P*ermE* knock-in | This study |
| pGWS1286 | pET28a with *gbn* coding sequence built-in, for $His_6$-Gbn fusion protein expression | This study |
| pGWS1300 | pUC19 harbouring partial *gbn* promoter region for EMSA experiment | This study |
| pGWS1451 | pUC19 harbouring random Gbn non-binding region for EMSA experiment | This study |
| pGWS1462 | pUC19 Harbouring partial *gbn* promoter region (-609 to +33) for TPM | This study |

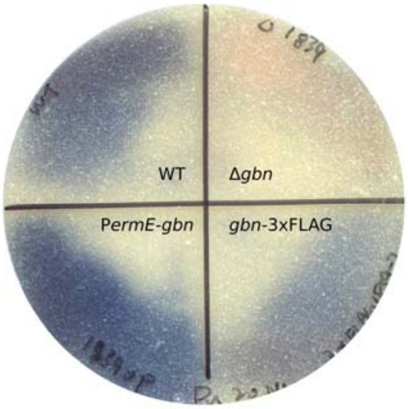


**Figure S1. SFM plate showing reduced production of the blue-pigmented actinorhodin (Act) in the *gbn* mutant. T**he *gbn* deletion strain (upper right) produces significantly less Act as the parental strain, while a strain over-expressing *gbn* (lower left) shows enhanced Act production. A strain expressing Gbn-3×FLAG (lower right) produces a similar amount of blue-pigmented Act as the parent (upper left).

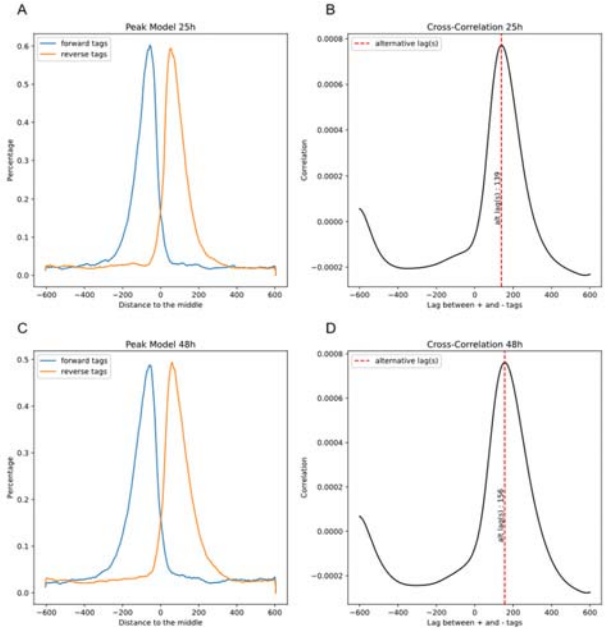


**Figure S2. MACS model of 25 h and 48 h ChIP-Seq data. (A, B)**, 25 h ChIP model, **(C, D)**, 48 h ChIP model. (A, C), 5' ends of strand-separated tags (reads) from a random sample of 1,000 model peaks, aligned by the centre of their Watson (forward) and Crick (reverse) peaks. (B, D), combined correlation peak from both strands, showing the distance (lag) of enriched tags to the centre of predicted binding region.

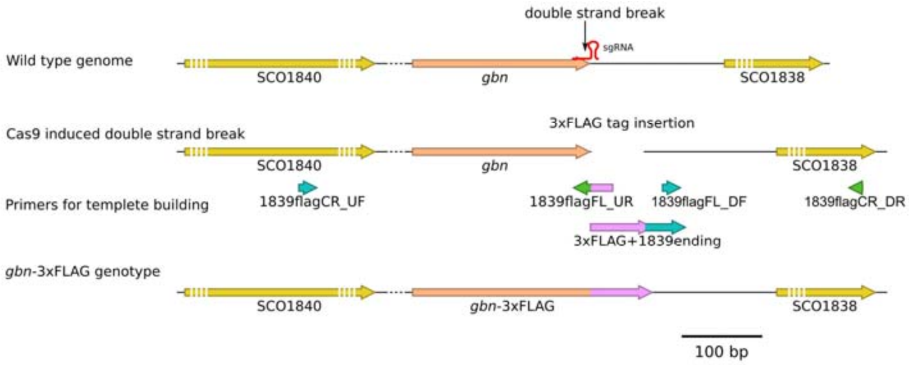


**Figure S3. Schematic of construct to allow 3×FLAG tag integration at the 3'-terminal of *gbn*.** Spacer sequence locates at the end of *gbn*, the sgRNA with this spacer guides the Cas9 protein to make a double-strand break after the stop codon. Templates for homology-directed repair (HDR) were made by cloning *gbn* and its upstream region from genome with additional connecting sequence, thus replacing the stop codon for connecting with 3×FLAG sequence. The downstream region was PCR-amplified from the genome proceeded by the sequence for a full 3×FLAG sequence. Then these two fragments were connected by overlap extension PCR.
